# Meclizine and metabotropic glutamate receptor agonists attenuate severe pain and primary sensory neuron Ca^2+^ activity in chemotherapy-induced peripheral neuropathy

**DOI:** 10.1101/2021.05.21.445188

**Authors:** John Shannonhouse, Matteo Bernabucci, Ruben Gomez, Hyeonwi Son, Yan Zhang, Hirotake Ishida, Yu Shin Kim

## Abstract

Chemotherapy-induced peripheral neuropathy (CIPN) affects about 68% of patients undergoing chemotherapy and causes severe neuropathic pain which is debilitating health problem and greatly reduces quality of life. Cisplatin is a commonly used platinum-based chemotherapeutic drug known to cause CIPN, possibly by causing oxidative stress damage to primary sensory neurons. Metabotropic glutamate receptors (mGluRs) are widely hypothesized to be involved in pain processing. Meclizine is an H1 histamine receptor antagonist which is known to have neuroprotective effects including anti-oxidative effect. Here, we used a mouse model of cisplatin-induced CIPN to test agonists of mGluR8 and group II mGluR as well as meclizine as interventions to reduce cisplatin-induced pain. We performed behavioral pain tests and *in vivo* entire DRG neurons Ca^2+^ imaging using genetically-encoded Ca^2+^ indicator, Pirt-GCaMP3 to monitor different drug interventions on a populational ensemble level. CIPN induced increased spontaneous Ca^2+^ activity in DRG neurons, increased Ca^2+^ transient amplitudes, and hyperresponses to mechanical, thermal, and chemical stimuli. We found mGluR8 agonist, DCPG, group II mGluR agonist, LY379268, and Histamine1 receptor antagonist, meclizine all significantly attenuated mechanical and thermal pain caused by CIPN. LY379268 and meclizine, but not DCPG, attenuated DRG neuronal Ca^2+^ activity elevated by CIPN. Furthermore, meclizine attenuated cisplatin-induced weight loss. These results suggest group II mGluR agonist, mGluR8 agonist, and meclizine are excellent candidates to study for mechanisms and new treatment option for CIPN.

## Introduction

Chemotherapy-induced peripheral neuropathy (CIPN) is a side effect of chemotherapy affecting about 68% of patients that causes severe neuropathic pain that persists for 6 months or longer in about 30% of patients who have undergone chemotherapy(1–5). Cisplatin is a platinum-based chemotherapeutic agent that inhibits tumor growth by crosslinking DNA nucleotides(6,7) and is widely used to treat cancer in various organs and body including lung, stomach, and head(5,8). There have been extensive studies on how to reduce neuropathic pain and its mechanisms at the level of individual cells, by using *in vitro* dorsal root ganglia (DRG) explants, and histology in animal models(2–4,9). However, CIPN has not been studied at *in vivo* entire DRG populational ensemble due to a lack of suitable tools and techniques, leaving an important gap in the understanding of CIPN mechanisms necessary to develop better therapeutics.

There are many non-mutually exclusive proposed mechanisms for CIPN caused by platinum-based chemotherapeutic drugs(9). These potential mechanisms including mitochondrial malfunction and cytoplasmic calcium unbalance(10), alteration of potassium channel subtype expression(11), upregulation of transient receptor potential vanilloid receptors(12,13), ER stress(14,15), primary sensory neuron senescence(16), and oxidative stress via mitochondrial disruption generating reactive oxygen species(ROS)(17–19). ROS produced by mitochondria are normally broken down by a pathway utilizing superoxide dismutase (SOD)(20). An SOD mimetic drug, calmangafodipir, reduced CIPN symptoms for oxaliplatin, a platinum-based drug(21). Interestingly, physiological changes during cisplatin-induced CIPN overlap extensively with known mechanisms of inflammatory pain(22–24).

Concomitant drug administration with cisplatin has reduced CIPN-induced neuropathic pain both in animal models and in clinical trials(19,21). So, *in vivo* DRG imaging may be particularly useful to study how concomitant drug administration with cisplatin modifies cisplatin’s effects on primary sensory neurons. All mGluRs except mGluR6 are found on central and/or peripheral neurons known to be involved in nociception and are known to affect pain(9,25–30). In particular, inflammatory pain is attenuated by group II mGluRs (mGluR2 and mGluR3) and group III mGluRs (mGlurR4, mGluR7, and mGluR8)(27,30,31) and enhanced or modulated by group I mGluRs (mGluR1 and mGluR5)(27,30–35). Thus, metabotropic glutamate receptors (mGluRs), especially group II and III, are promising therapeutic targets for CIPN-induced neuropathic pain(27,30).

Meclizine is another attractive candidate for reducing CIPN from platinum-based drugs. Meclizine is an H1 receptor antagonist and a pregnane X receptor agonist in mice shown to be neuroprotective in animal models of diverse conditions including Huntington’s(36), Parkinson’s(37), and hypoxia(38). Meclizine protects cultured dorsal root ganglia (DRG) primary sensory neurons from cisplatin-induced damage by enhancing pentose phosphate pathway, enhancing NADPH production, and improving clearance of DNA nucleotides damaged by cisplatin(18). Furthermore, meclizine shifts retinal ganglion cells towards glycolysis metabolism pathway but away from mitochondrial respiratory pathway(39), suggesting it could be protective against cisplatin-induced oxidative stress and mitochondrial toxicity.

In this study, we used mechanical and thermal pain behavior assays and confocal microscopy imaging with the fluorescent genetically-encoded calcium indicator, Pirt-GCaMP3 to investigate cisplatin-based CIPN-induced neuropathic pain in *in vivo* entire DRG primary sensory neurons on a populational ensemble level. We found cisplatin induced mechanical and thermal hypersensitivity in pain behavior assays, increased levels of spontaneous Ca^2+^ activity in DRG primary sensory neurons, and increased Ca^2+^ responses to mechanical, thermal, and chemical stimuli. mGluR8 agonist DCPG, group II mGluRs (mGluR2 and mGluR3) agonist LY379268, and histamine H1 receptor antagonist meclizine each attenuated cisplatin-induced pain in behavior assays. Furthermore, LY379268 and meclizine strongly attenuated spontaneous calcium activity in *in vivo* DRG neurons. Finally, meclizine significantly reduced cisplatin-induced weight loss.

## Materials and Methods

### Animals

All experiments were performed in compliance with Institutional Animal Care and Use Committee at University of Texas Health Science Center at San Antonio (UTHSA) and in accordance with National Institutes of Health and American Association for Accreditation of Laboratory Animal Care. Pirt-GCaMP3 mice(40,41) in a C57BL/6J background or C57BL/6J mice (Jackson Laboratory, Bar Harbor, ME, USA) were used in all experiments. Pirt-GCaMP3 mice used in experiments were all heterozygous. Both males and females were used. All animals were at least 8 weeks old, were kept in a 14/10 hour light/ dark cycle and had *ad libitum* access to food and water.

### Drugs and drug treatment

Cisplatin, LY378268, and DCPG were purchased from Abcam (Cambridge, MA, USA). Meclizine was purchased from Tocris (Minneapolis, MN, USA). All compounds were dissolved in 0.9% sterile saline and injected i.p. unless otherwise specified. Isoflurane was purchased from the Piramal Group (Mumbai, India). Pentobarbital was purchased from Diamondback Drugs (Scottsdale, AZ, USA). Cisplatin and saline vehicle injections were performed on days 1, 3, 5, and 7. In experiments using multiple rounds of cisplatin injection, cisplatin was also injected on days 12, 14, 16, and 18 except in the LY379268 experiments where the second round of cisplatin injections were given on days 17, 19, 21, and 23. When DCPG was concomitantly administered to animals receiving cisplatin, injections were given 30 minutes prior to cisplatin. When meclizine was concomitantly administered with cisplatin, injections were given 3 hours prior to cisplatin. When LY379268 was administered, it was given 2, 4, 6, and 8 days after the last cisplatin injection.

### Behavioral Tests

#### von Frey mechanical test

Mice were placed in a 4.5cm x 5cm x 10cm transparent container on a metal mesh and allowed to habituate for at least 30 minutes prior to testing. Each mouse was tested at a specific force 8 times in order to determine the lowest force required to elicit a paw withdrawal response more than 50% of the time.

#### Hot plate thermal test

Mice were placed in a 4.5cm x 5cm x 10cm transparent container on a temperature controlled hot plate set to 45°C. Latency to acute nocifensive behavior was determined by onset of hindpaw lifts, licking, jumping, or flinching.

### DRG exposure surgery

L5 DRG exposure surgery and imaging was performed as previously described(41). Mice were anesthetized with pentobarbital (40-50mg/kg). After deep anesthesia was achieved, the animal’s back was shaved and the shaved skin was aseptically prepared by cleaning alcohol and iodine pad and ophthalmic ointment was applied to keep the eyes moist (Lacrilube, Allergan Pharmaceuticals, Irvine, CA, USA). Mice were kept on a heating pad and monitored with a rectal thermometer to keep body temperature at 37±0.5°C.

Dorsal laminectomy was performed in the L4-L6 area. A 2cm long midline incision was made in the lower back around the lumbar enlargement area. Paravertebral muscles were dissected away to expose the lower lumbar enlargement area and the bones were cleaned. Small rongeurs were used to remove the surface aspect of the L5 DRG transverse process bone near the vertebra to expose the DRG without disrupting the neurons, axons, and other cells in the DRG.

### *In vivo* DRG GCaMP Ca^2+^ imaging

*In vivo* entire DRG GCaMP Ca^2+^ imaging was performed during the 1-6 hours following exposure surgery. Mice were kept on a heating pad and monitored with a rectal thermometer to keep body temperature at 37±0.5°C. Mice were laid abdomen down on a custom-designed imaging stage. Movement from breathing, heart beats, etc. was minimized by holding the head in a custom designed holder with an anesthesia/ gas mask and custom designed vertebral clamps. Continuous anesthesia was maintained by 1%-2% isoflurane in pure oxygen.

For imaging and analysis, the stage was fixed under a single photon confocal microscope (Carl Zeiss AG, Oberkochen, Germany). Raw image stacks (512×512 to 1024×1024 pixels in the x-y plane and 20-30μm voxel depth) were converted into time lapse movies (it took ~6.5 to 7 seconds to produce a single frame) and analyzed using Zeiss Zen 3.1 Blue Edition software (Carl Zeiss AG, Oberkochen, Germany).

Putative responding cells were identified by visual observation of raw image time lapse movies. Ca^2+^ transient intensities were calculated by ΔF / F_0_ = (F_t_ – F_0_) / F_0_ where F_t_ is the pixel intensity in a region of interest (ROI) at the time point of interest and F_0_ is the baseline intensity determined by averaging the intensities of the first 2-6 frames of the ROI in the experiment. For calcium responses to stimuli that produced multiple transients, only the first peak was analyzed. Cells showing calcium transients before the stimulus were assumed to be spontaneously active. Each peak of spontaneous Ca^2+^ transients was analyzed individually.

Stimuli were applied carefully to not cause movement during imaging. Press (SMALGO Algometer, Bioseb Instruments, Vitrolles, France) and von Frey filaments were applied directly to the hindpaw ipsilateral to the DRG being imaged for 15-20 seconds. Thermal stimuli (0°C and 45°C) were applied by immersing the hindpaw in water at a specific temperature for 15-20 seconds. Stimuli were applied after 35-40 seconds of baseline imaging. LY379268 during imaging was applied by topically pipetting to DRG neurons. Small and large brush bristles were 5 mm and 40 mm, respectively.

### Statistical Analysis

Statistics were performed on GraphPad Prism 9.0.1. Ca^2+^ transient intensities and numbers of activated cells showing spontaneous Ca^2+^ activity were analyzed by Student’s t-Test or One-Way or Two-Way ANOVA followed by a post-hoc Tukey’s Test or post-hoc Dunnett’s Test. Activated cell number counts following stimuli were analyzed by Two-Way ANOVA. Body weight was analyzed by two-way ANOVA and either Tukey’s or Dunnett’s post-hoc Tests.

## Results

### Cisplatin induces mechanical and thermal hyperalgesia, increases spontaneous Ca^2+^ activity in the DRG neurons, and increases the sensitization of DRG neurons to mechanical and thermal stimuli

In order to determine the effects of cisplatin on DRG Ca^2+^ signaling, animals were injected with cisplatin (3.5mg/kg, i.p.) every other day for 4 total injections(42). Compared to saline-treated controls, cisplatin-injected animals developed mechanical hyperalgesia after the second injection of cisplatin (Fig. 1A) and thermal hyperalgesia after the fourth cisplatin injection (Fig. 1B). Ca^2+^ imaging experiments were performed as previously described(41). Animals were anesthetized and DRGs were surgically exposed and imaged. In cisplatin-injected mice, we found robust increases in the number of cells with spontaneous Ca^2+^ oscillation and total spontaneous Ca^2+^ activity (steady-state high Ca^2+^ levels and Ca^2+^ oscillation) but not number of cells with steady-state high Ca^2+^ activity after cisplatin injections compared to saline-injected mice (Table 1). The number of DRG neurons responding to press (100g, 300g and 600g) and 45°C stimuli were increased compared to saline controls (Table 1).

**Figure 1.**
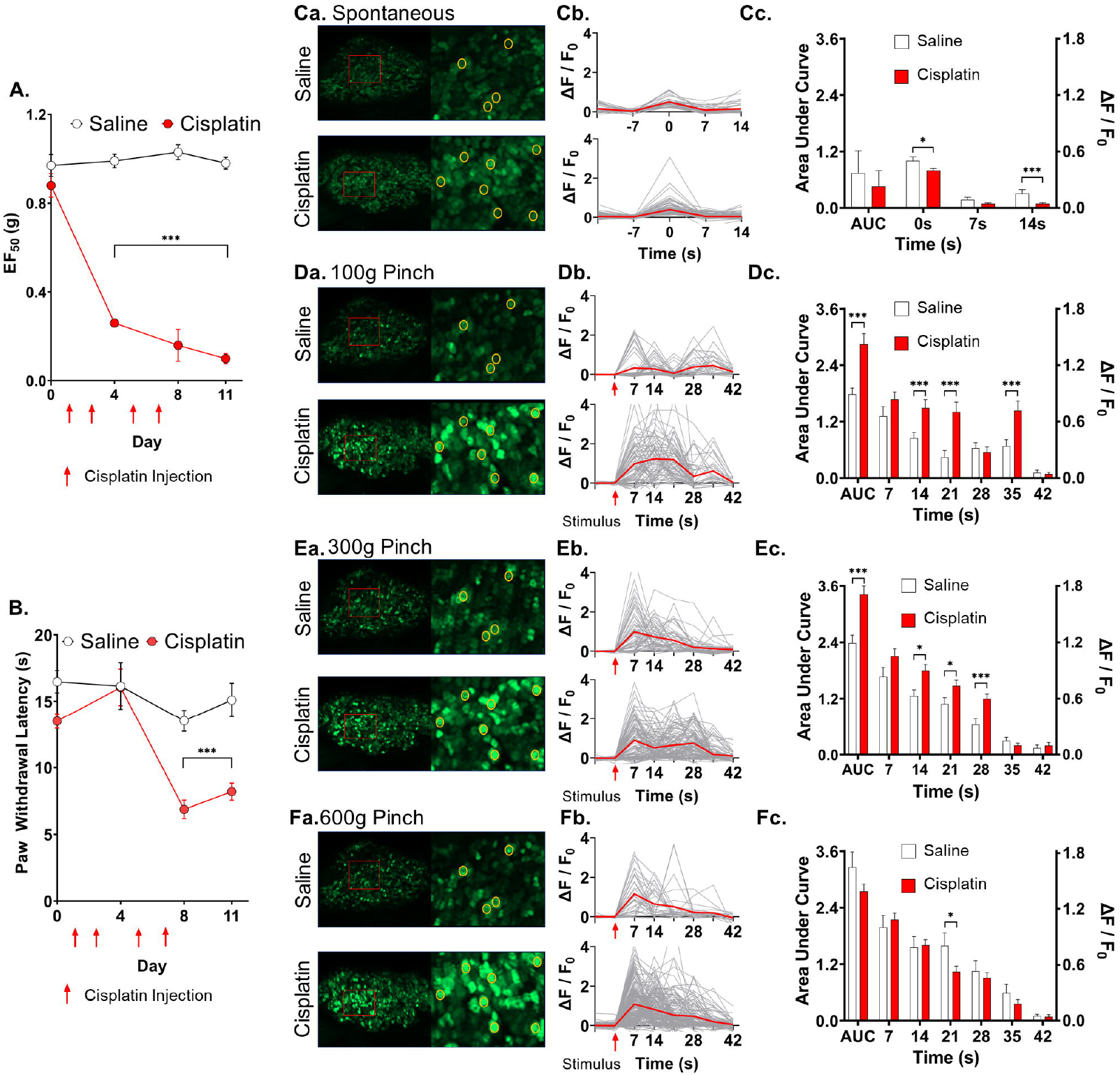
Cisplatin treatment causes increased sensitivity to pain and increased Ca^2+^ responses to press stimuli. (**A**) Mechanical pain sensitivity tests using von Frey filaments were performed on days 0, 4, and 8. Red arrows indicate saline or cisplatin injection (days 1, 3, 5, and 7). Mechanical sensitivity is plotted as 50% withdraw threshold in grams. (**B**) Thermal pain sensitivity tests using hot plate assay were performed on days 0, 4, 8, and 11. Red arrows indicate saline or cisplatin injection (days 1, 3, 5, and 7). Thermal sensitivity is plotted as paw withdraw latency in seconds. (**Ca**) Representative images of DRG neurons spontaneous Ca^2+^ activity in saline- and cisplatin-injected animals by maximum intensity Z-projections. Entire DRGs are shown on left. Area outlined by red boxes are magnified on the right. Yellow circles indicate spontaneously Ca^2+^ activated neurons and later panels in different stimuli. (**Cb**) Graphs of DRG neurons spontaneous Ca^2+^ transients are shown with maximum fluorescence intensity set at 0s. Y axis is expressed as ΔF / F_0_ fluorescence intensity. (**Cc**) Mean fluorescence intensities of spontaneous Ca^2+^ transients are shown as a function of time. Left Y axis shows area under the curve in arbitrary unit. Right Y axis is expressed in ΔF / F_0_ fluorescence intensity. (**Da**, **Ea**, **Fa**) Representative images of DRG neurons Ca^2+^ activity in response to press stimulus (100g, 300g, and 600g, respectively) of saline- and cisplatin-injected animals. The same entire DRGs from panel **Ca** are shown on left. Area outlined by red boxes are magnified on the right. Yellow circles indicate Ca^2+^ activated neurons in different stimuli. (**Db**, **Eb**,**Fb**) Graphs of Ca^2+^ transients in response to press stimuli. Red arrow indicates start of press stimuli. Y axis is expressed in ΔF / F_0_ fluorescence intensity. (**Dc**, **Ec**, **Fc**) Mean fluorescence intensities of Ca^2+^ transients in response to press stimuli. Left Y axis shows area under the curve in arbitrary unit. Right Y axis is expressed in ΔF / F_0_ fluorescence intensity for each frame. Comparisons were performed with two-way ANOVA followed by Tukey’s post hoc tests. (Saline vs. cisplatin * p < 0.05, ** p < 0. 01, *** p < 0.001)

**Table 1.** Number of cells activated spontaneously or following stimuli. Neurons with spontaneous activity in saline control animals, and animals-treated with cisplatin, cisplatin + DCPG, and cisplatin + meclizine are shown. One-way ANOVAs with Dunnett’s post-hoc t-tests comparing all groups to cisplatin-treated animals and DCPG + cisplatin and meclizine + cisplatin treatment to cisplatin-treated animals were performed. Stimulus response comparisons between saline and cisplatin-treated animals are shown. Stimulus responses were compared by two-way ANOVA (treatment x stimulus) and post-hoc Tukey tests were performed for each stimulus. Compared to saline * P<0.05, ** p<0.01, ***p<0.001; compared to cisplatin # p<0.05, ## p<0.01

| **Spontaneous Activity** |  |  |  |  |
| --- | --- | --- | --- | --- |
|  | Saline (activated cell number) | Cisplatin | Cisplatin + DCPG | Cisplatin + Meclizine |
| Spontaneous Ca<sup>2+</sup> Oscillation | 5.60 ± 1.69, n = 5, p<0.05<sup>#</sup> | 17.50 ± 3.32, n = 6 | 18.80 ± 9.57, n = 5, p>0.05 | 7.40 ± 3.93, n = 5, p>0.05 |
| Steady-state high Ca<sup>2+</sup> | 6.00 ± 2.76, n = 5, p>0.05 | 16.33 ± 4.94, n = 6 | 9.00 ± 3.35, n = 5, p>0.05 | 7.00 ± 1.79, n = 5, p>0.05 |
| Total Cells with Ca<sup>2+</sup> Activity | 11.60 ± 3.72, n = 5, P<0.01<sup>##</sup> | 33.83 ± 5.25, n = 6 | 27.80 ± 8.33, n = 5, p>0.05 | 14.40 ± 2.93, n = 5, p<0.05<sup>#</sup> |
| **Stimulus Responses** |  |  |  |  |
|  | Saline (activated cell number) | Cisplatin |  |  |
| Press 100g | 43.75 ± 5.76, n = 4 | 73.50 ± 12.25, n = 4, p<0.05* |  |  |
| Press 300g | 60.25 ± 4.71, n = 4 | 120.75 ± 18.44, n = 4, p< 0.05* |  |  |
| Press 600g | 90.33 ± 16.26, n = 3 | 129.33 ± 3.67, n = 3, p<0.05* |  |  |
| Small Brush | 2.29 ± 0.87, n = 7 | 3.75 ± 0.25, n = 4, p>0.05 |  |  |
| Large Brush | 4.17 ± 1.42, n = 6 | 6.20 ± 0.86, n = 5, p>0.05 |  |  |
| 0°C | 14.60 ± 1.78, n = 5 | 19.25 ± 0.63, n = 4, p>0.05 |  |  |
| 45°C | 26.75 ± 1.60, n = 4 | 66.67 ± 9.82, n = 3, p<0.01** |  |  |
| Von Frey 0.07g | 1.83 ± 0.54, n = 6 | 3.20 ± 0.92, n = 5, p>0.05 |  |  |
| Von Frey 10g | 17.83 ± 2.66, n = 5 | 26.80 ± 3.97, n = 5, p>0.05 |  |  |

Stimulus-induced Ca^2+^ transients were monitored in saline- and cisplatin-treated animals. The number of cells showing spontaneous Ca^2+^ activity and Ca^2+^ transients in response to stimuli was higher in DRGs of cisplatin-treated animals than saline-treated animals (F_1,47_ = 49.21, p < 0.001 by two-way ANOVA). Analysis of spontaneous and press-induced Ca^2+^ transients shows that spontaneous Ca^2+^ transients in the DRG of saline-treated control took longer to return to baseline than cisplatin-treated animals (Fig. 1Cb, Cc). At 100g and 300g press, cisplatin-treated animals had greater area under the curve and were slower return to baseline than in saline-treated animals (Fig. 1Db, Dc, Eb, Ec). Ca^2+^ transients by 600g press were not statistically different between saline- and cisplatin-treated animals (Fig. 1F). Compared to Ca^2+^ transients in DRG neurons of saline-treated mice, cisplatin treatment increased area under the curve of Ca^2+^ transients in response to small brush stimulus and decreased area under the curve of Ca^2+^ transients in response to paw immersion in 45°C water (Fig. 2Ab, Ac, Db, Dc). Compared to saline-treated controls, more Ca^2+^ transients were delayed in cisplatin-treated animals in response to large brush stimulus (Fig. 2Bc). Amplitude of Ca^2+^ transients in saline-treated animals in response to paw immersion in 45°C water was significantly higher compared to cisplatin-treated animals (Fig. 2Dc). Paw immersion in 0°C water and paw stimulus with 0.07g and 10g von Frey did not show statistically detectable differences between Ca^2+^ transients of two groups (Fig. 2Cb, Cc, Eb, Ec, Fb, Fc).

**Figure 2.**
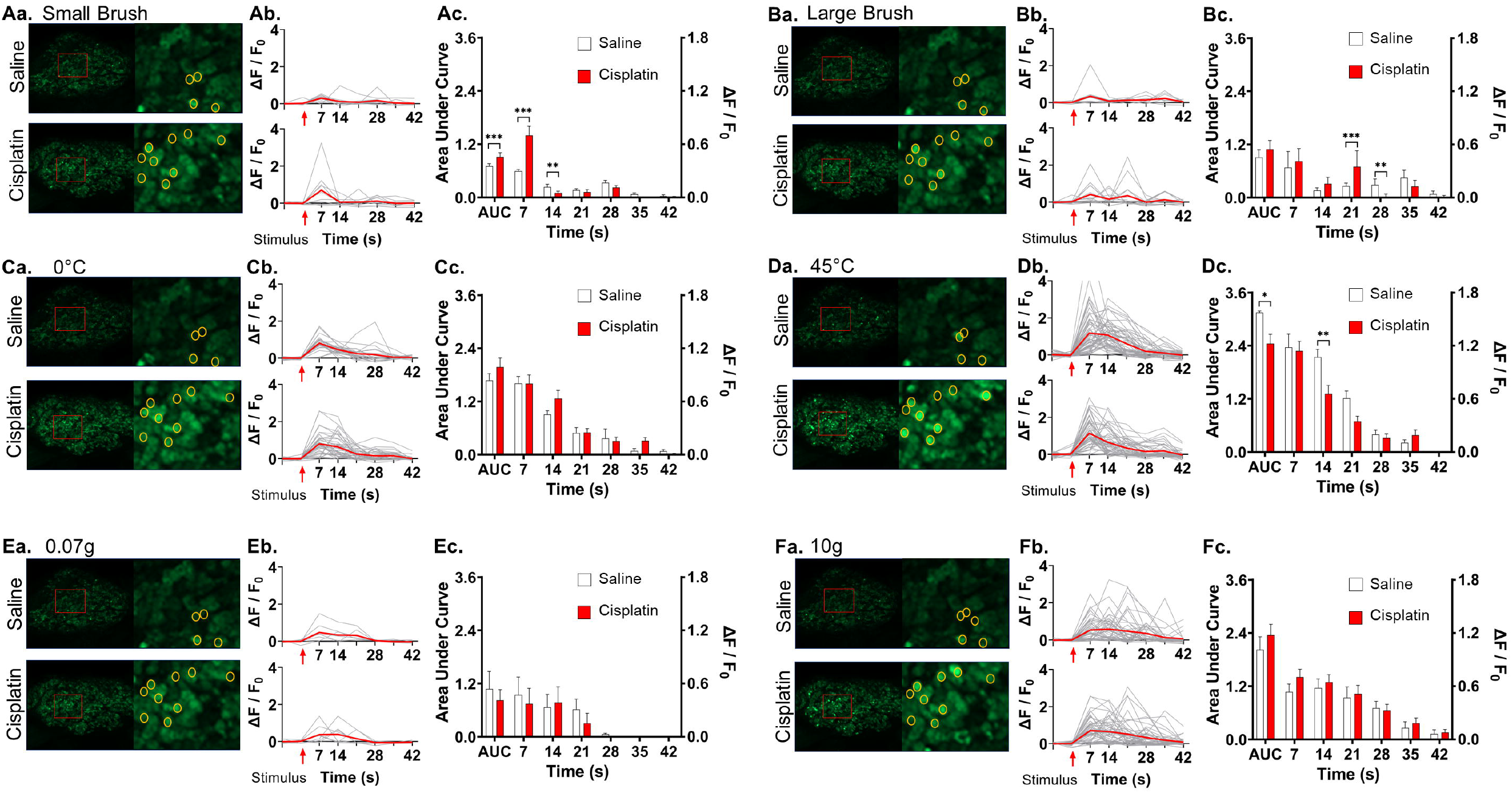
Cisplatin treatment affects DRG neurons Ca^2+^ activity in response to light to strong stimuli. (**Aa**, **Ba**, **Ca**, **Da**, **Ea**, **Fa**) Representative images of DRG neurons Ca^2+^ activity in response to stimuli (small brush (**Aa**), large brush (**Ba**), 0°C cold (**Ca**), 45°C hot (**Da**), 0.07g von Frey (**Ea**), and 10g von Frey (**Fa**), respectively) of saline- and cisplatin-injected animals in maximum intensity Z-projections. The same entire DRGs are shown in each panel on left. Area outlined by red boxes are magnified on the right. Yellow circles indicate DRG neurons Ca^2+^ activity in different stimuli. (**Ab**, **Bb**, **Cb**, **Db**, **Eb**, **Fb**) Graphs of Ca^2+^ transients in response to various stimuli. Red arrow indicates start of each stimulus. Y axis is expressed in ΔF / F_0_ fluorescence intensity. (**Ac**, **Bc**, **Cc**, **Dc**, **Ec**, **Fc**) Mean fluorescence intensities of Ca^2+^ transients in response to various stimuli. Left Y axis shows area under the curve in arbitrary unit. Right Y axis shows amplitude in ΔF / F_0_ fluorescence intensity for each frame. Comparisons were performed with two-way ANOVA followed by Tukey’s post hoc tests. (Saline vs. cisplatin * p < 0.05, ** p < 0.01, *** p < 0.001)

Relative proportions of small, medium, and large diameter neurons on spontaneously oscillating Ca^2+^ activity and responding to stimuli were indistinguishable between cisplatin-treated and saline-treated mice. Cell activation and its consequential Ca^2+^ transients by spontaneous activity and most stimuli predominantly happened in smaller diameter neurons (<20μm) followed by medium diameter neurons (20-25μm) with relatively few large diameter neurons (>25μm). However, stronger mechanical stimuli (10 gram von Frey, 300g and 600g press) activated a higher proportion of large diameter neurons than other stimuli (Supplementary Tables S1, S2).

### Cisplatin-induced hyperalgesia was attenuated by concomitant administration of mGluR8 agonist, DCPG

Since mGluR8 agonists are known to attenuate some kinds of inflammatory pain(26,43), we tested mGluR8 agonist, DCPG in the cisplatin-induced pain model by injecting DCPG 30 minutes before each cisplatin injection. DCPG robustly attenuated cisplatin-induced mechanical and thermal hyperalgesia (Fig. 3A, B). However, neither the number of activated neurons showing spontaneous Ca^2+^ activity nor the number of activated neurons in response to 45°C, 0.07g von Frey, or 10g von Frey stimuli were statistically distinguishable between cisplatin and cisplatin with DCPG-treated animals (Table 1, Table 2). The number of activated neurons following 0°C cold stimulus was increased in cisplatin with DCPG-treated animals compared to cisplatin alone animals (Table 2). DCPG slowed spontaneous Ca^2+^ transients’ decay time (return to baseline) (Fig. 3Cb, Cc). DCPG also increases the neuronal number of delayed Ca^2+^ transients in response to 0°C cold (Fig. 3Db, Dc) but decreased area under the curve of Ca^2+^ transients in response to 10g von Frey stimulus (Fig. 3Gb, Gc).

**Figure 3.**
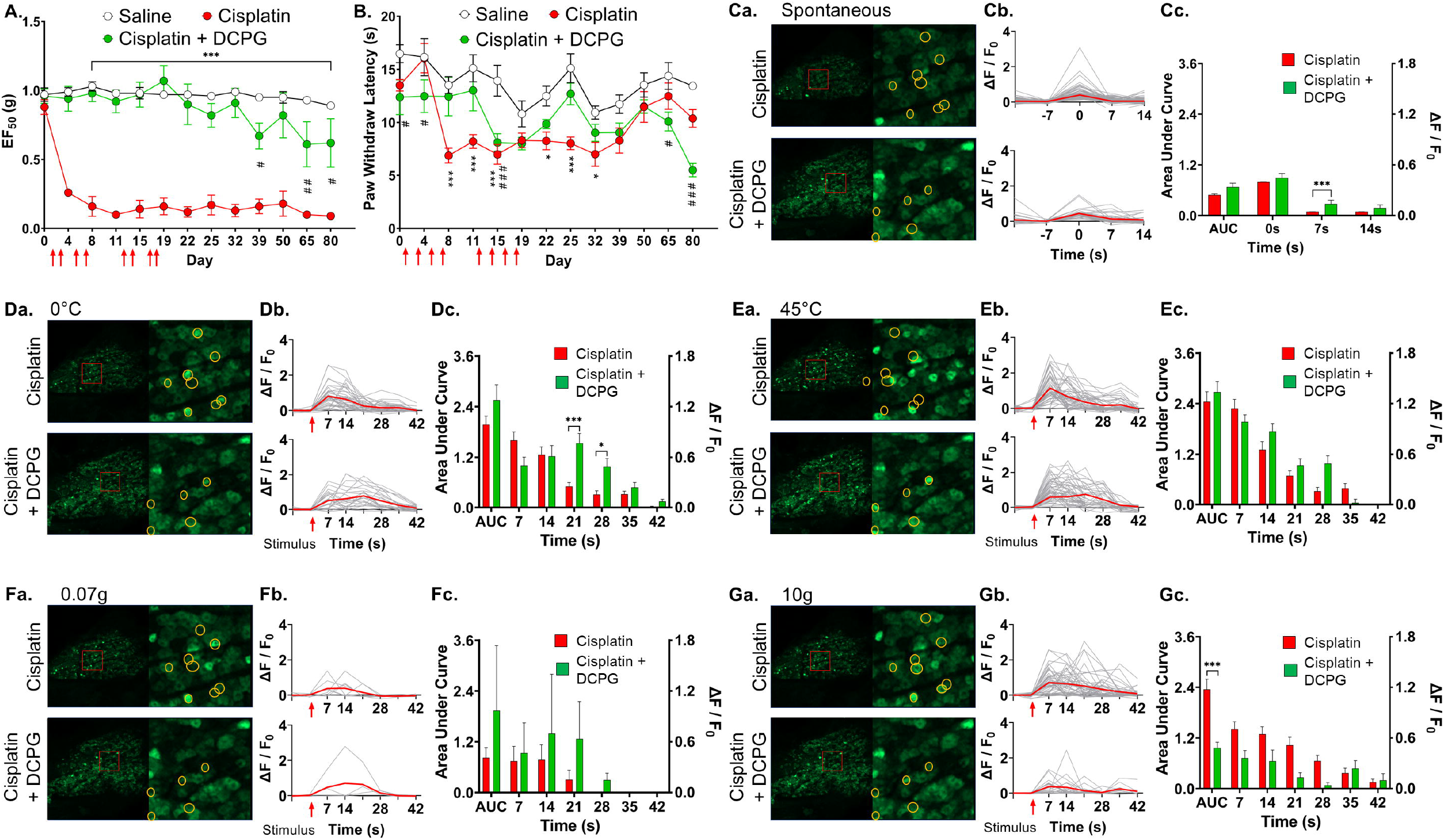
Metabotropic glutamate receptor 8 agonist, DCPG reduces mechanical and thermal pain sensitivity in cisplatin-treated animals and affects Ca^2+^ activity in response to some stimuli. (**A**) Mechanical pain sensitivity tests using von Frey filaments were performed on every 3-15 days up to 80 days. Red arrows indicate drug injection (days 1, 3, 5, 7 and 12, 14, 16, 18) of saline or DCPG followed 30 minutes later by cisplatin. Mechanical sensitivity is plotted as 50% withdraw threshold in grams. (**B**) Thermal pain sensitivity tests using hot plate assay were performed on every 3-15 days up to 80 days. Red arrows indicate drug injection (days 1, 3, 5, 7 and 12, 14, 16, 18) of saline or DCPG followed 30 minutes later by cisplatin. Thermal sensitivity is plotted as paw withdraw latency in seconds. (**Ca**) Representative images of DRG neurons spontaneous Ca^2+^ activity in cisplatin-, or cisplatin + DCPG injected animals in maximum intensity Z-projections. Entire DRGs are shown on left. Area outlined by red boxes are magnified on the right. Yellow circles indicate Ca^2+^ activating neurons by various stimuli in later panels. (**Cb**) Graphs of DRG neurons spontaneous Ca^2+^ transients are shown with maximum fluorescence intensity set at 0s. Y axis is expressed in ΔF / F_0_ fluorescence intensity. (**Cc**) Mean fluorescence intensities of DRG neurons spontaneous Ca^2+^ transients as a function of time frame. Left Y axis shows area under the curve in arbitrary unit. Right Y axis indicates amplitude in ΔF / F_0_ fluorescence intensity for each frame. (**Da**, **Ea**, **Fa**, **Ga**) Representative images of maximum intensity Z-projections of Ca^2+^ transients in response to different stimuli (0°C cold (**Da**), 45°C hot (**Ea**), 0.07g von Frey (**Fa**), and 10g von Frey (**Ga**), respectively) of cisplatin- and cisplatin + DCPG-injected animals. The same entire DRGs in panel **Ca** are shown on left. Area outlined by red boxes are magnified on the right. Yellow circles indicate DRG neurons activated by different stimuli. (**Db**, **Eb**, **Fb**, **Gb**) Graphs of DRG neurons Ca^2+^ transients in response to different stimuli (0°C cold (**Db**), 45°C hot (**Eb**), 0.07g von Frey (**Fb**), and 10g von Frey (**Gb**), respectively). Red arrow indicates start of each stimulus. Y axis is expressed in ΔF / F0 fluorescence intensity. (**Dc**, **Ec**, **Fc**, **Gc**) Mean fluorescence intensities of DRG neurons Ca^2+^ transients in response to various stimuli. Left Y axis shows area under the curve in arbitrary unit. Right Y axis shows amplitude in ΔF / F_0_ fluorescence intensity for each frame. All comparisons were performed using two-way ANOVA followed by Tukey’s post hoc tests. (For behavior: Saline vs. cisplatin * p < 0.05, ** p < 0.01, *** p < 0.001; cisplatin vs. cisplatin + DCPG # p < 0.05, ## p < 0.01, ### p < 0.001, for Ca^2+^ transients: cisplatin vs. cisplatin + DCPG * p < 0.05, ** p < 0.01, *** p < 0.001)

**Table 2.** Number of cells activated following thermal and mechanical stimuli. Comparisons of stimulus responses of saline, cisplatin, cisplatin + DCPG, and cisplatin + meclizine to hindpaw immersion in 0°C water, 45°C water, and 0.07g, 10g von Frey to the hindpaw. Stimulus responses were compared by two-way ANOVA (treatment x stimulus) followed by post-hoc Dunnett’s test compared to cisplatin-treated animals. Compared to cisplatin-treated # p<0.05, ## p<0.01, ### p<0.001

| ***Cells showing Ca<sup>2+</sup> transients in response to stimuli*** |  |  |  |  |
| --- | --- | --- | --- | --- |
| Stimulus | Saline | Cisplatin | Cisplatin + DCPG | Cisplatin + Meclizine |
| 0°C | 14.60 ± 1.78, n = 5, p>0.05 | 19.25 ± 0.63, n = 4 | 37.33 ± 4.37, n = 3, p<0.01<sup>##</sup> | 16.20 ± 3.65, n = 5 |
| 45°C | 26.75 ± 1.60, n = 4, p<0.001<sup>###</sup> | 66.67 ± 9.82, n = 3 | 50.75 ± 6.34, n = 4, p>0.05 | 22.40 ± 5.63, n = 5, p<0.001<sup>###</sup> |
| Von Frey 0.07g | 1.83 ± 0.54, n = 6, p>0.05 | 3.20 ± 0.92, n = 5 | 2.00 ± 0.45, n = 5, p>0.05 | 4.20 ± 1.28, n = 5, p>0.05 |
| Von Frey 10g | 17.83 ± 2.66, n = 5, p>0.05 | 26.80 ± 3.97, n = 5 | 38.75 ± 7.12, n = 4, p>0.05 | 22.50 ± 4.87, n = 4, p>0.05 |

### Cisplatin-induced hyperalgesia and the number of spontaneous Ca^2+^ activated cells were attenuated by group II mGluR agonist, LY379268

Group II metabotropic glutamate receptors are strong candidates for analgesic targets for inflammatory pain(27), so, we tested mGlu2/3 receptor agonist, LY379268 for effects on cisplatin-induced pain. LY379268 was injected (3mg/kg, i.p.) for four days following cisplatin injections (3.5mg/kg, i.p.). We found that LY379268 caused animals to rapidly recover from cisplatin-induced mechanical hyperalgesia but had no detectable effect on animals after they had recovered from cisplatin-induced mechanical hyperalgesia (Fig. 4A). LY379268 also prevented sustained cisplatin-induced thermal hyperalgesia (Fig. 4B). The number of spontaneous Ca^2+^ activated neurons per DRG was decreased in a pairwise test when LY379268 was topically applied to DRGs of cisplatin-treated mice during imaging sessions (Fig. 4C). Ca^2+^ transients in spontaneously activated neurons before and 10-15 minutes after LY379268 application were compared, but no statistically detectable differences were found in area under the curve, curve shape (rise or decay time change), or the number of Ca^2+^ oscillations in pairwise tests (Fig. 4Db, Dc, Fig. 4E).

**Figure 4.**
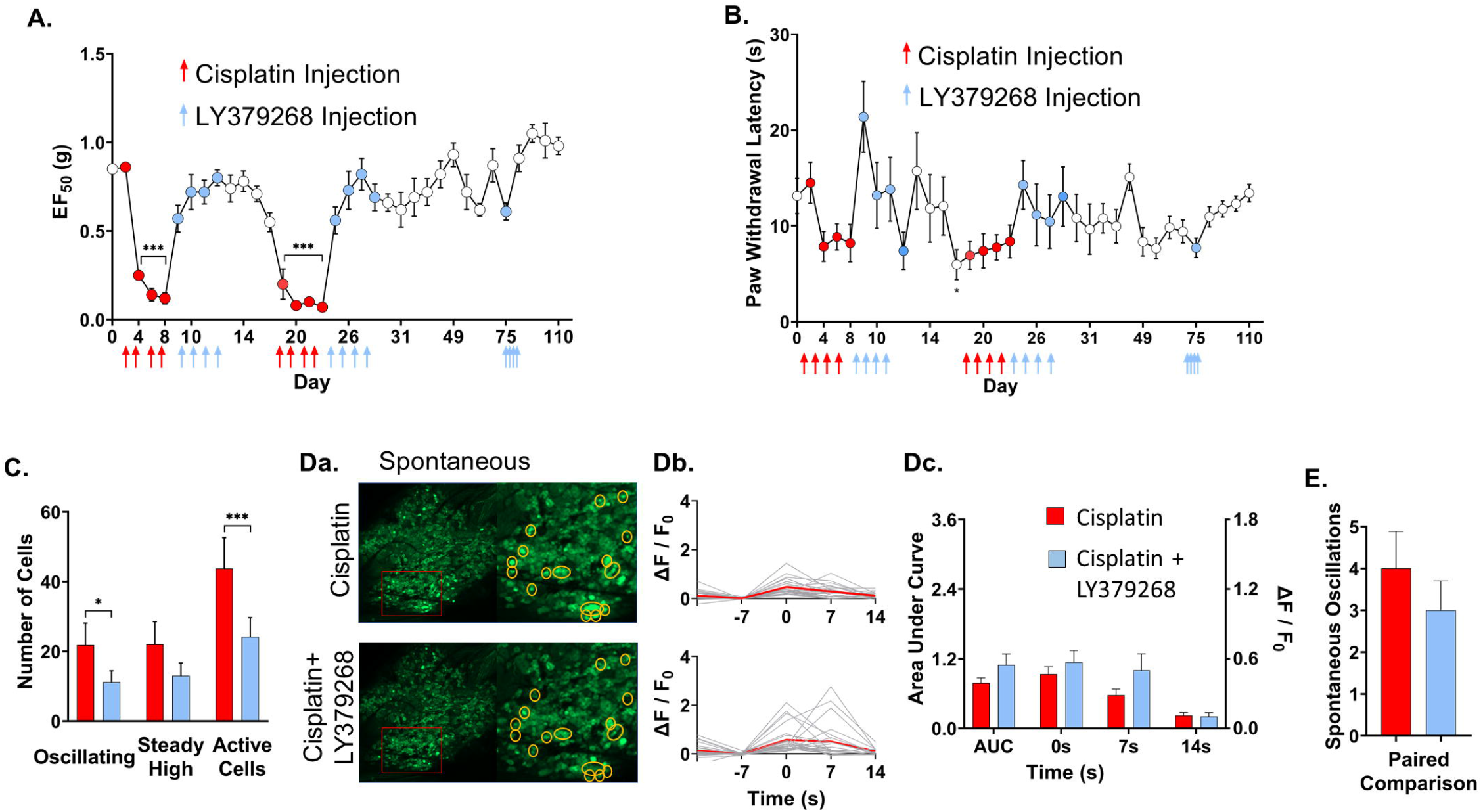
Metabotropic glutamate receptors 2/3 agonist, LY379268 reduces mechanical and thermal pain sensitivity in cisplatin-treated animals and reduces DRG neurons spontaneous Ca^2+^ activity. (**A**-**B**) Mechanical pain sensitivity tests using von Frey filaments and thermal pain sensitivity tests using hot plate were performed every on every 2-15 days. Red arrows indicate drug injection of cisplatin (1, 3, 5, 7, 17, 19, 21, and 23). Blue arrows indicate days of LY379268 injection (daily 9-12, daily 25-28, and daily 75-79). Mechanical sensitivity is plotted as 50% withdraw threshold in grams. Thermal sensitivity is plotted as paw withdraw latency in seconds. Comparisons to baseline (day 0) were made using one-way ANOVA followed by Dunnett’s post hoc comparisons. (**C**) Number of DRG neurons with Ca^2+^ oscillation, Steady-state high, and total numbers of DRG neurons Ca^2+^ activity before and 15 minutes after topical application of LY379238 to the DRG. Comparisons are paired t-tests. (**Da**) Representative images of DRG neurons spontaneous Ca^2+^ activity before and after LY379268 administration. Entire DRGs are shown on left. Area outlined by red boxes are magnified on the right. Yellow circles indicate spontaneously Ca^2+^ activating neurons. (**Db**) Spontaneous Ca^2+^ transients of DRG neurons before and after LY379268 application. Maximum fluorescence intensity set at 0s. Y axis is expressed in ΔF / F_0_ fluorescence intensity. (**Dc**) Mean fluorescence intensities of DRG neurons spontaneous Ca^2+^ transients before and after LY379268 application. Left Y axis shows area under the curve in arbitrary unit. Right Y axis shows amplitude in ΔF / F_0_ fluorescence intensity for each frame. (**E**) Mean number of DRG neurons Ca^2+^ transients before and after LY379268 application. Comparison is by paired t-test. (For behavior: each time point vs. baseline Day 0 * p < 0.05, ** p < 0.01, *** p < 0.001, for Ca^2+^ activity: cisplatin vs. LY379268 * p < 0.05, ** p < 0.01, *** p < 0.001)

### Cisplatin-induced mechanical hyperalgesia, spontaneous DRG neuronal Ca^2+^ activity, and Ca^2+^ response to 45°C hot stimulus were all attenuated by histamine receptor 1 antagonist, meclizine

Since meclizine has a protective role against DNA damage by cisplatin(18), we tested meclizine in cisplatin-treated animals for its effects on behavior and Ca^2+^ activity. Meclizine was injected (16mg/kg, i.p.) three hours prior to cisplatin injection (3.5mg/kg, i.p.) 4 times over 8 days. Meclizine robustly reduced cisplatin-induced mechanical hypersensitivity (Fig. 5A). Meclizine also robustly attenuated spontaneous Ca^2+^ activity in cisplatin-treated animals, bringing the number of neurons with spontaneous Ca^2+^ activity (Ca^2+^ oscillation and steady-state high Ca^2+^ together) down to numbers seen in saline-treated animals (Table 1 and Fig. 5B), and reduced the number of Ca^2+^ activated neurons in response to 45°C water (thermal stimulus) (Fig. 5Da). In addition, meclizine treatment significantly increased area under the curve of Ca^2+^ transients and decay time (return to baseline) in response to 0°C water (cold stimulus) (Fig. 5Cb, Cc). In contrast, meclizine reduced area under the curve of Ca^2+^ transients in response to 45°C water (Fig. 5Db, Dc) and 10g von Frey (Fig. 5Fb, Fc). Meclizine caused no statistically detectable change on Ca^2+^ transients in response to 0.07g von Frey (Fig. 5Eb, Ec).

**Figure 5.**
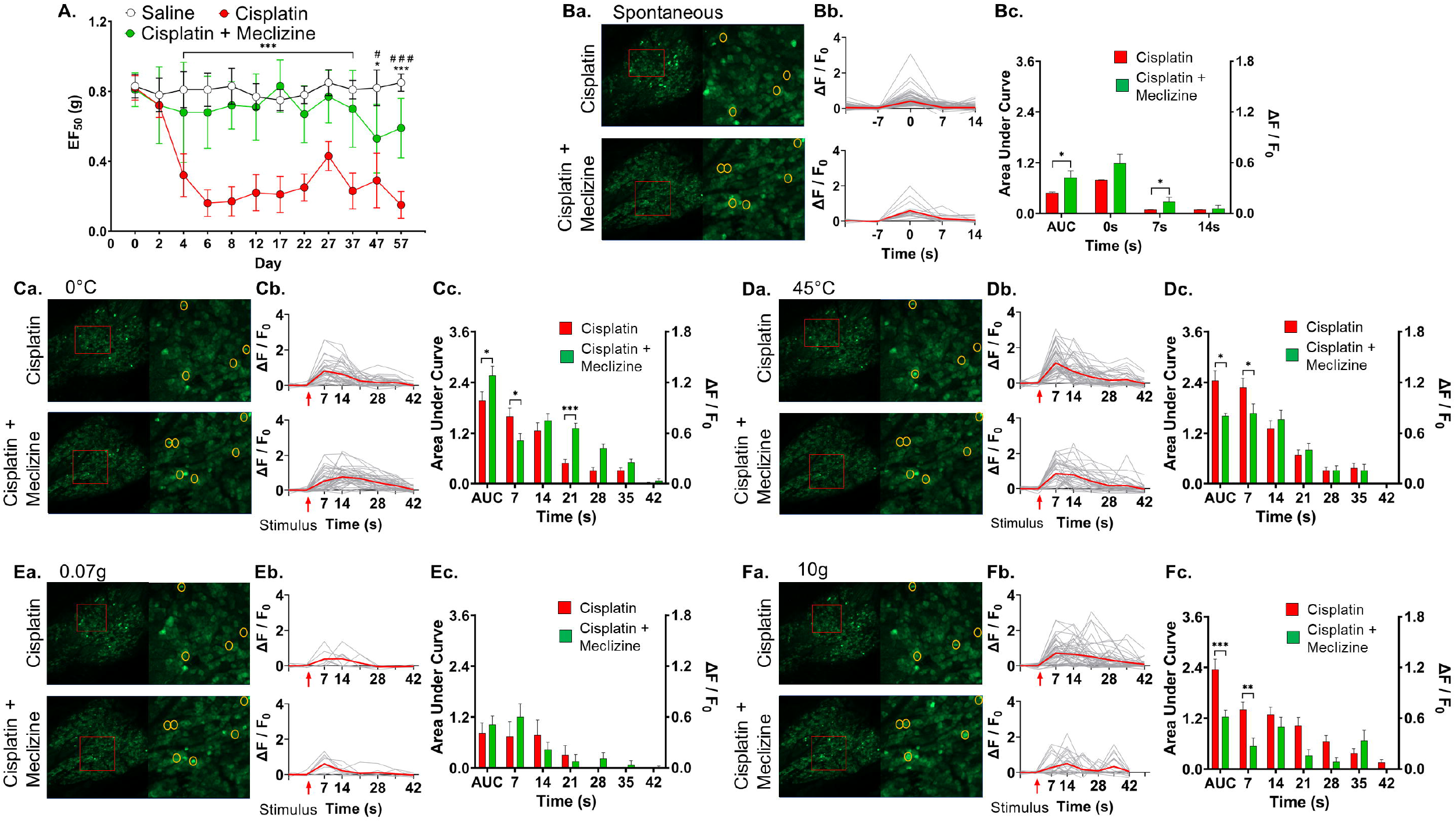
Meclizine attenuates cisplatin-induced mechanical pain sensitivity and reduces DRG neurons Ca^2+^ activity in response to thermal and mechanical stimuli. (**A**) Mechanical pain sensitivity tests using von Frey filaments were performed on every 2-10 days. Red arrows indicate saline, cisplatin or cisplatin + meclizine injection (days 1, 3, 5, and 7). Meclizine or saline vehicle was injected 3 hours prior to cisplatin injection. Mechanical sensitivity is plotted as 50% withdraw threshold in grams. (**Ba**) Representative images of DRG neurons spontaneous Ca^2+^ activity in cisplatin-, or cisplatin + meclizine injected animals. Entire DRGs are shown on left. Area outlined by red boxes are magnified on the right. Yellow circles indicate spontaneously activating neurons. (**Bb**) Graphs of DRG neurons spontaneous Ca^2+^ transients are shown with maximum fluorescence intensity set at 0s. Y axis is expressed in ΔF / F_0_ fluorescence intensity. (**Bc**) Mean fluorescence intensities of spontaneous Ca^2+^ transients as a function of time frame. Left Y axis shows area under the curve in arbitrary unit. Right Y axis shows amplitude in ΔF / F_0_ fluorescence intensity for each frame. (**Ca**, **Da**, **Ea**, **Fa**) Representative images of DRG neurons Ca^2+^ activity in response to stimuli (0°C cold (**Ca**), 45°C hot (**Da**), 0.07g von Frey (**Ea**), and 10g von Frey (**Fa**), respectively) of cisplatin- and cisplatin + meclizine-injected animals. The same entire DRGs as in panel **Ba** are shown on left. Area outlined by red boxes are magnified on the right. Yellow circles indicate Ca^2+^ activating neurons with different stimuli. (**Cb**, **Db**, **Eb**, **Fb**) Graphs of DRG neurons Ca^2+^ transients in response to various stimuli. Red arrow indicates start of each stimulus. Y axis shows ΔF / F_0_ fluorescence intensity. (**Cc**, **Dc**, **Ec**, **Fc**) Mean fluorescence intensities of DRG neurons Ca^2+^ transients in response to various stimuli. Left Y axis shows area under the curve in arbitrary unit. Right Y axis shows amplitude in ΔF / F_0_ fluorescence intensity for each frame. All comparisons were performed using two-way ANOVA followed by Tukey’s post hoc tests. (For behavior: Saline vs. cisplatin * p < 0.05, ** p < 0.01, *** p < 0.001, cisplatin vs. cisplatin + meclizine # p < 0.05, ## p < 0.01, ### p < 0.001, for Ca^2+^ transients: cisplatin vs. cisplatin + meclizine * p < 0.05, ** p < 0.01, *** p < 0.001)

### Cisplatin-induced weight loss was attenuated by meclizine but unaffected by DCPG and LY379268

Cachexia, a condition characterized by weight loss, muscle loss, and adipose loss, is associated with cancer and greatly increases cancer mortality(44), and is exacerbated by nearly every form of chemotherapy(45,46). Since drugs were administered by proportion of animal weight, animal weight was monitored over the time course of experiments. Cisplatin treatment caused rapid, persistent weight loss (Fig. 6A, B, C). DCPG had no positive effect on body weight loss (Fig. 6A). Mean body weight of DCPG-treated animals took 32 days after the last cisplatin injection to return to starting mean body weight and LY379268 took 26 days (Fig. 6A, B). Surprisingly, however, animals that received concomitant meclizine administration rapidly recovered mean body weight. Meclizine-treated animals recovered all mean weight loss by 10 days after the last injection. Meclizine-treated animals’ mean weight began decreasing by 50 days after their last injection (Fig. 6C).

**Figure 6.**
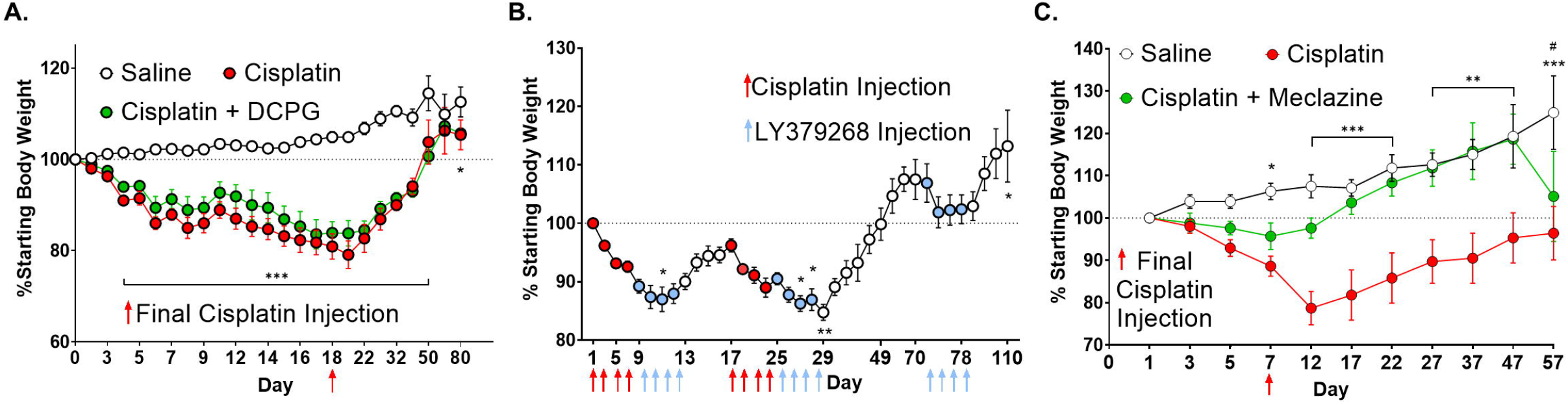
Cisplatin treatment results in significant body weight loss in mice which is attenuated by meclizine treatment but not by DCPG or LY379268 treatment. (A) Graph of body weight over the course of mechanical and thermal sensitivity experiments with saline-, cisplatin-, and cisplatin + DCPG-injected animals (Fig. 3). Y axis is expressed in % starting body weight. Red arrow indicates the day of final cisplatin injection. Comparisons were performed using two-way ANOVA followed by Tukey’s post hoc tests. (B) Graph of body weight over the course of cisplatin- and LY379268-injected animals (Fig.4). Red arrow indicates when cisplatin injection is performed. Blue arrow indicates when LY379268 injection is performed. Comparisons were performed using one-way ANOVA with post hoc Dunnett’s tests. (C) Graph of body weight over the course of mechanical and thermal sensitivity experiments with saline-, cisplatin-, and cisplatin + meclizine-injected animals (Fig.5). Red arrow indicates the day of final cisplatin injection. Comparisons were performed using two-way ANOVA followed by Tukey’s post hoc tests. (Saline vs. cisplatin * p < 0.05, ** p < 0.01, *** p < 0.001; saline vs. cisplatin + meclizine # p < 0.05)

## Discussion

CIPN is painful, debilitating, and complex phenomenon occurring in patients treated with many chemotherapeutic drugs. Chemotherapeutic drugs act through different molecular and cellular mechanisms, suggesting causes of CIPN are similarly diverse (1–5). This in turn suggests the best ways to treat CIPN may vary by the kind of chemotherapeutic drug used. The DRG is a site of peripheral sensory neuropathy. Therefore, it is important to be able to study the effects of specific chemotherapeutic agents and specific therapies on peripheral neurons and how they affect responses to different mechanical, thermal, and chemical stimuli. However, until recently direct *in vivo* observation of the DRG was very challenging due to limited tools and techniques. *In vivo* entire DRG neurons Ca^2+^ imaging is a valuable tool that allows study of neurons firing and activity at a populational level as an ensemble and allows for detection and study of physiologically important phenomena that would be very challenging to detect using other methods such as electrophysiological recordings(41).

Prior research has shown cisplatin treatment induces spontaneous activity and hyperexcitability in dissociated primary sensory neurons *in vitro* (47). In this study, we demonstrated *in vivo* entire DRG neurons Ca^2+^ activity and Ca^2+^ transients in response to several mechanical and thermal stimuli were increased by cisplatin treatment as an animal model of CIPN. Concomitant meclizine administration attenuated CIPN mechanical pain hypersensitivity and concomitant administration of DCPG or LY379268 attenuated CIPN mechanical and thermal pain hypersensitivity. Our results also showed two of three drugs that attenuated CIPN pain hypersensitivity also attenuated the number of spontaneous Ca^2+^ activated neurons. Meclizine attenuated Ca^2+^ transients in response to 45°C. Furthermore, concomitant meclizine administration greatly reduced cisplatin-induced weight loss.

DCPG’s effect on cisplatin-induced CIPN pain appears to be outside of the DRG neurons. mGluR8 is present in most DRG(48,49) and trigeminal ganglia(50) neurons. However, its role in CIPN has never been investigated. Peripheral administration of mGluR8 agonists can attenuate thermal and mechanical hyperalgesia caused by carrageenan, formalin, and capsaicin(26,30,48). DCPG can modulate pain centrally, as well. Infusion of DCPG into the periaqueductal gray (PAG) was sufficient to reduce nociceptive responses and thermal and mechanical hypersensitivity caused by formalin and carrageenan injection and PAG infusion of an antagonist of Group III mGluRs reduced the analgesic effect of peripheral DCPG, showing mGluR8 can act fully or partially through a central anti-nociceptive pathway(26). Administering mGluR8 agonist DCPG 30 minutes prior to cisplatin produced a striking reduction of cisplatin-induced mechanical hypersensitivity and reduced thermal hypersensitivity, yet DCPG-treated animals were not distinguishable in terms of numbers of Ca^2+^ activated cells or numbers of activated cells responding to any stimulus analyzed in this study except 0°C, where DCPG increased the number of responding neurons. While there were some subtle differences in Ca^2+^ transients, DCPG failed to produce the striking effects on DRG neurons Ca^2+^ activity seen with LY379268 and meclizine. Since DCPG administration was systemic, it is not possible to distinguish central and peripheral effects. The lack of striking decreases in numbers of Ca^2+^ activated neurons spontaneously or from stimuli combined with DCPG’s known effects in the PAG suggests DCPG’s attenuation of cisplatin-induced mechanical and thermal hypersensitivity is central rather than peripheral or that DCPG works peripherally through a mechanism that does not affect DRG neurons Ca^2+^ activity. mGluR8 may be a target for CIPN pain even though it does not seem to act peripherally.

Group II metabotropic glutamate receptor agonists attenuate hyperalgesia and allodynia from a broad range of causes(27,51–54) and antagonists of group II receptors aggravate hyperalgesia or allodynia, block analgesia, and/or prolong recovery time from these conditions(28,29,38) as well as increase the number of action potentials in primary sensory neurons in response to capsaicin or heat(55). Group II mGluRs (mGluR2/3) are broadly expressed in primary sensory neurons(49,50,54). Our data show that in this cisplatin-induced CIPN model, mGluR2/3 agonist LY379268 decreased the number of DRG neurons spontaneous Ca^2+^ activity, reduces mechanical hyperalgesia, and blocks extended thermal hypersensitivity. Mechanical and thermal sensitivity assays were based on systemic administration, so it is not possible to determine to what extent the effect was peripheral or central. However, reduced numbers of DRG Ca^2+^ activated neurons were a direct and acute effect. Acute reduction of spontaneous Ca^2+^ activity when LY379268 was applied directly to the DRG neurons during imaging suggests analgesic efficacy is at least partially peripheral. Other studies have shown peripheral mGluR2 or mGluR2/3 agonists can block inflammatory hyperalgesia(28,56,57). Side effects remain a major concern for group II mGluR agonists due to convulsions observed in preclinical animal models(58). Limiting a mGluR2/3 drug to the periphery may reduce side effects(59–61). Our results combined with earlier research suggest a mGluR2/3 agonist with poor blood brain barrier penetration could treat CIPN pain without disrupting mGluR2/3 signaling in the central nervous system.

Because meclizine treatment was systemic, it is impossible to distinguish a central from peripheral effect. However, there are good reasons to believe meclizine acts through mitochondrial repair and/or reducing oxidative stress. Cisplatin binds mitochondrial DNA in DRG neurons, inhibits mitochondrial transcription and replication, and causes mitochondrial DNA degradation(62). Cisplatin directly inhibits mitochondrial respiration system *in vivo* and reduces transcription and increases degradation of mitochondrial RNA(63). Meclizine improves clearance of damaged mitochondrial DNA(18), by shifting mitochondrial respiration towards glycolysis metabolism(37,38). Meclizine also maintains an elevated ratio of reduced: oxidized glutathione and NADPH(18,38). Therefore, meclizine has multiple mechanisms to protect neurons from oxidative damage. Reactive nitrogen species (ROS) and oxidative stress are known to activate and/or potentiate a variety of transient receptor potential channels (64,65), which may partially depolarize neurons thus, making them more excitable and resulting in high spontaneous Ca^2+^ activity and decreasing neurons firing threshold in response to stimuli. ROS also affect potassium and sodium channels(66,67) as well as damaging a wide range of other molecules in the cell(68). By improving redox state, enhancing DNA repair, and shifting primary sensory neuron metabolism away from mitochondrial respiration and towards glycolysis metabolism, meclizine may attenuate hyperexcitability through multiple known mechanisms (Fig. 7) and therefore reduce hypersensitivity to painful stimuli. Meclizine is also an approved drug for use in humans making it an excellent candidate to treat CIPN.

**Figure 7.**
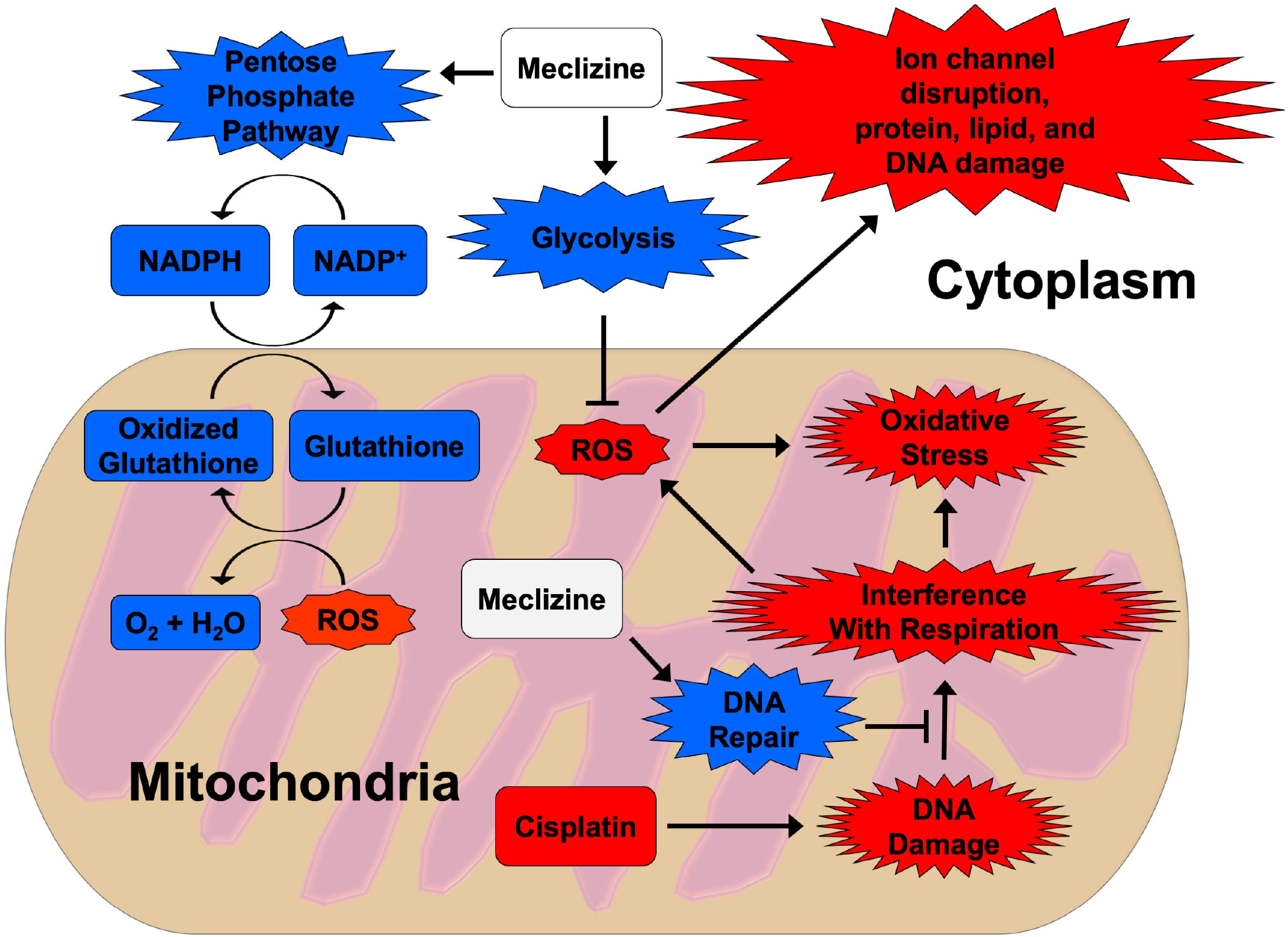
In an oxidative stress damage model of cisplatin-induced CIPN, meclizine can prevent spontaneous Ca^2+^ activity and reduce sensitization of sensory neurons through various mechanisms. Cisplatin interferes with mitochondrial respiration and causes mitochondrial DNA damage. This in turn produces reactive oxygen species (ROS) which cause oxidative stress and can directly activate some transient receptor potential channels. Oxidative stress and transient receptor potential channels activation cause primary sensory neuron hyperexcitability, leading to high levels of spontaneous Ca^2+^ activity and hypersensitivity to stimuli. Meclizine has three known mechanisms to combat oxidative stress damage from cisplatin: (1) meclizine promotes glycolysis and lactate production in neurons, reducing mitochondrial respiration and resultant ROS production. (2) Meclizine stimulates repair of cisplatin-mediated DNA damage, resulting in healthier mitochondria. Meclizine stimulates the pentose phosphate pathway to reduce NADP to NADPH which in turn reduces glutathione (GSH) allowing GSH to reduce mitochondrial ROS.

One of the most common side effects of platinum-based chemotherapeutic drugs is weight loss(44–46). There have been many mechanisms proposed for how cisplatin and other platinum-based chemotherapeutic drugs cause weight loss(69–76). Meclizine reduced both cisplatin-induced hypersensitivity to mechanical pain and attenuated cisplatin-induced weight loss. There were no detectable effects of DCPG or Ly379268 on body weight of cisplatin-treated animals. Meclizine is commonly used for treatment of nausea and vomiting during pregnancy(77) and motion sickness(78). Cisplatin induces nausea in humans(79). Meclizine is known to be a strong antagonist of H1 histamine receptor, a weak antagonist of muscarinic acetylcholine receptor(80), and a weak antagonist of constitutive androstane receptor in mice(81). Histamine suppresses food intake in the central nervous system(82,83) and histaminergic neurons play a large role in food intake, metabolism, weight gain or loss, and a foraging-related locomotor activities(83). Mice lacking the muscarinic 3 acetylcholine receptor eat less and weight less than wild type mice(84,85) and stimulation of hypothalamic muscarinic acetylcholine receptors stimulates food intake(86). Meclizine could potentially attenuate weight loss through antagonism of anorexigenic H1 histamine receptors or muscarinic acetylcholine receptors.

We have used a cisplatin-based CIPN model to study *in vivo* DRG neurons at a populational level. We have found two (LY379268 and meclizine) of three drug treatments that attenuated CIPN-induced pain hypersensitivity and reduced spontaneous DRG neuron Ca^2+^ activity. Of the two treatments tested in response to a variety of stimuli, meclizine decreased CIPN-induced DRG neuron hypersensitivity to stimuli. The mGluR8 agonist treatment that didn’t reduce spontaneous Ca^2+^ activity is known to be able to reduce pain hypersensitivity centrally in some inflammatory pain models. These results demonstrate *in vivo* DRG neuron Ca^2+^ imaging can be used to study mechanisms of CIPN and peripherally-acting candidates for therapeutic intervention.

## Supplemental Data

**Supplemental Table S1.** Comparisons of cell diameter of DRG neurons responding to press stimuli in saline-treated controls and cisplatin-treated animals. Most activated neurons were small (<20μm) or medium (20μm-25μm) diameter. Strong mechanical stimuli (300g and 600g press) activated a higher proportion of large diameter (>25μm) neurons. Proportions of small v. medium v. large diameter neurons were not statistically distinguishable between groups.

| ***Press Responses*** |  |  |
| --- | --- | --- |
| ***100g press: number of activated cells (proportion of activated cells)*** |  |  |
| Diameter | Saline | Cisplatin |
| Small (<20µm) | 22.00 ± 4.51, n = 3 (0.559 ± 0.084) | 46.33 ± 7.75, n = 3 (0.676 ± 0.038) |
| Medium (20-25µm) | 12.67 ± 3.53, n = 3 (0.336 ± 0.087) | 17.67 ± 2.40, n = 3 (0.252 ± 0.002) |
| Large (>25µm) | 4.00 ± 0.58, n = 3 (0.105 ± 0.020) | 4.33 ± 2.19, n = 3 (0.072 ± 0.037) |
| ***300g press: number of activated cells (proportion of activated cells)*** |  |  |
| Small | 40.33 ± 5.90, n = 3 (0.534 ± 0.067) | 52.00 ± 12.12, n = 3 (0.519 ± 0.127) |
| Medium | 23.67 ± 2.03, n = 3 (0.316 ± 0.034) | 29.67 ± 6.12, n = 3 (0.294 ± 0.187) |
| Large | 11.33 ± 2.33, n = 3 (0.150 ± 0.036) | 19.00 ± 8.62, n = 3 (0.187 ± 0.079) |
| ***600 g press: number of activated cells (proportion of activated cells)*** |  |  |
| Small | 49.00 ± 15.72, n = 3 (0.512 ± 0.075) | 64.00 ± 7.77, n = 3 (0.495 ± 0.056) |
| Medium | 26.33 ± 2.96, n = 3 (0.314 ± 0.056) | 41.67 ± 2.03, n = 3 (0.320 ± 0.021) |
| Large | 15.00 ± 2.52, n = 3 (0.174 ± 0.031) | 23.67 ± 5.36, n = 3 (0.184 ± 0.043) |

**Supplemental Table S2.**
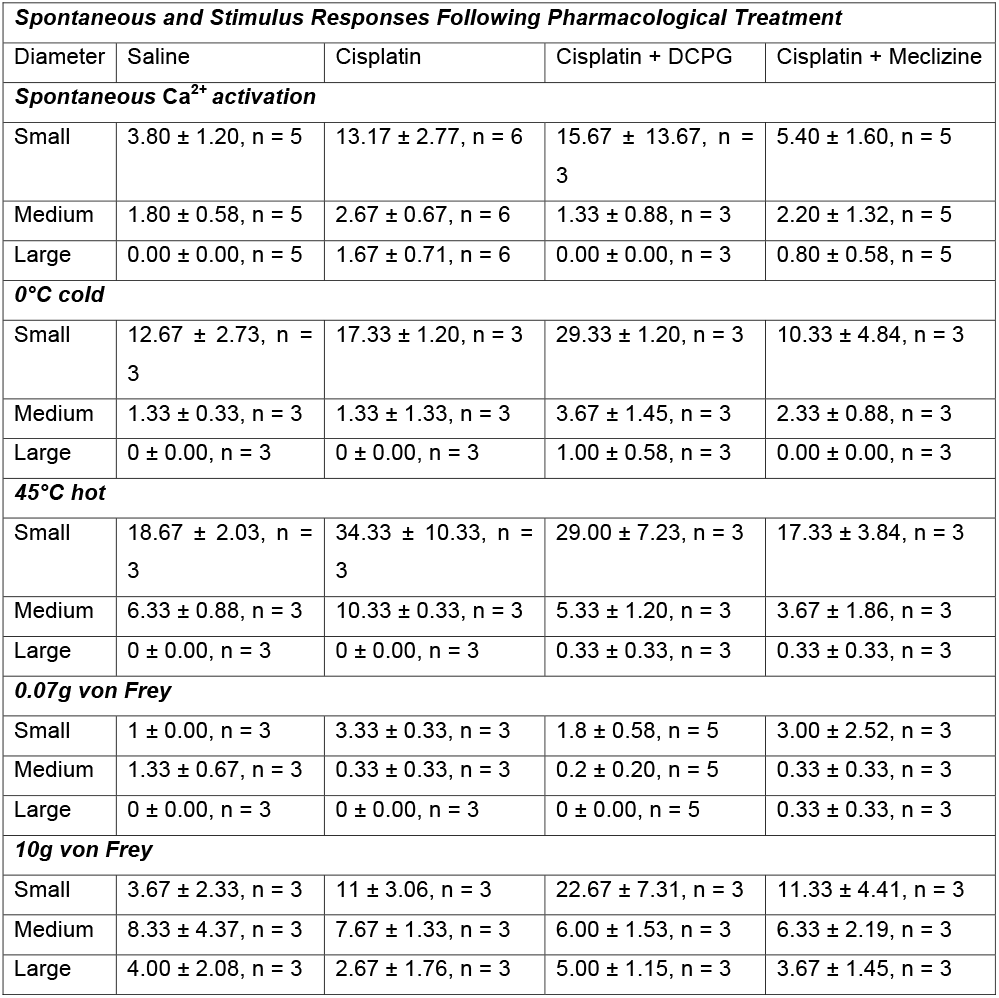
Comparisons of cell diameter of DRG neurons responding to different stimuli in saline-treated controls and cisplatin-treated, cisplatin + DCPG, and cisplatin + meclizine treated animals. Most activated neurons were small (<20μm) or medium (20μm-25μm) diameter. Strong mechanical stimulus (10g von Frey) tended to activate a higher proportion of large diameter (>25μm) neurons.

## Supplemental Movie Legends

**Supplemental Movie 1:** Cisplatin dramatically increases the number of DRG neurons responding to mechanical stimulus. Representative movies generated from microscope images compare DRGs during 100g press between saline-treated and cisplatin-treated animals. In response to a non-noxious press (100g), an average of 43.75 cells produces Ca^2+^ transients in saline-treated animals (**1A**) and an average of 73.50 cells in cisplatin-treated animals (**1B**).

**Supplemental Movie 2:** Acute treatment with group II mGluR agonist, LY379268 decreases spontaneous Ca^2+^ activity in cisplatin-treated animals. Representative movies generated from microscope images compare DRG spontaneous Ca^2+^ activity in cisplatin-treated animals before and after LY379268 treatment. In cisplatin-treated animals that were subsequently treated with LY379268, an average of 43.8 neurons produced spontaneous Ca^2+^ activity (Ca^2+^ oscillations or steady-state high Ca^2+^ activity) (**2A**). After LY379268 treatment, an average of 24.2 neurons produced spontaneous Ca^2+^ activity (**2B**).

**Supplemental Movie 3:** Meclizine dramatically decreases the number of DRG neurons responding to paw immersion in 45°C water in cisplatin-treated animals. In animals treated with cisplatin, an average of 66.67 neurons showed Ca^2+^ transients in response to 45°C hot stimulus (**3A**). Treatment with meclizine 3 hours prior to cisplatin injection decreased the average number of activated DRG neurons to 22.40 (**3B**).

## Acknowledgements

This work was supported by National Institutes of Health Grant (R01DE026677 to Y.S.K.), UTHSCSA startup fund (Y.S.K.), and a Rising STAR Award from University of Texas system (Y.S.K.).

## Author Contributions

M.B. and Y.S.K contributed to study design with assistance from J.S., R.G., Y.S.K. contributed to data interpretation and manuscript revision. M.B. performed experiments with assistance from J.S., H.I., H.S., Y.Z., and J.S. drafted the paper. R.G. and J.S. maintained, set up mating, took care of mice, and performed genotyping. R.G. assisted with GCaMP imaging work. Y.S.K. supervised all aspects of the project and wrote the paper.

## Conflicts of Interest

Authors declare no conflicts of interest and no financial conflicts.

## Notes

### Competing Interest Statement

The authors have declared no competing interest.

